# Integrated Human-Virus Metabolic Modelling Predicts Host-Based Antiviral Targets Against Chikungunya, Dengue and Zika Viruses

**DOI:** 10.1101/165605

**Authors:** Sean Aller, Andrew E. Scott, Mitali Sarkar-Tyson, Orkun S. Soyer

## Abstract

Current and reoccurring viral epidemic outbreaks such as those caused by Zika virus illustrate the need for rapid development of antivirals. Such development would be immensely facilitated by computational approaches that can provide experimentally testable predictions for possible antiviral strategies. A key factor that has not been considered fully to date in the study of antiviral targets is the high dependence of viruses to their host metabolism for reproduction. Here, we focus on this dependence and develop a stoichiometric, genome-scale metabolic model that integrates human macrophage cell metabolism with the biochemical demands arising from virus production. Focusing this approach to currently epidemic viruses Chikungunya, Dengue and Zika, we find that each virus causes specific alterations in the host metabolic flux towards fulfilling their individual biochemical demands as predicted by their genome and capsid structure. Subsequent analysis of this integrated model allows us to predict a set of host reactions, which when constrained can inhibit virus production. We show that this prediction recovers most of the known targets of existing antiviral drugs, while highlighting a set of hitherto unexplored reactions with either broad or virus specific antiviral potential. Thus, this computational approach allows rapid generation of experimentally testable hypotheses for novel antiviral targets within a host.

**SIGNIFICANCE STATEMENT:** A key challenge in combatting any new and emerging virus outbreaks is rapid drug development. In particular, generation of experimentally testable hypotheses through computational approaches is mostly lacking. Here, we address this gap by developing host-virus metabolic models for three viruses that cause current (or previously) epidemic viral outbreaks. We develop viral biomass functions using information from their genomes and physical structure, and incorporate these within a genome-scale metabolic model of human macrophage cells. The resulting integrated model allows us to predict host reactions, which when blocked, stop the system from attaining optimal viral production. These predictions recover currently known antiviral targets within human cells, and highlight a set of new reactions that are hitherto not explored for antiviral capacity.

## INTRODUCTION

Rapid development of antiviral drugs for emerging and re-emerging viruses, such as the Zika virus, remains a significant challenge (1, 2). Given that virus production within a host is intertwined with host immune response and metabolism (3), it is suggested that novel development of antivirals should take into account host processes (4, 5). Indeed, viruses are entirely dependent on their hosts’ cellular resources for their replication. This is highlighted by observed variations in virus production levels correlating with cell-to-cell variance in growth rate and phase (6), as well as virus infection leading to changes in host metabolism (7). In particular, virus infection leads to significant metabolic alterations in the host, in some cases resulting in up to 3-fold increase in glycolysis rates (7-9) and changes in ATP production rates (6). This observation can be seen as an emergent property of the combined host-virus metabolic system and could be related to changes in host cellular demands arising from viral production (10, 11). More specifically, alterations in host metabolism upon infection can be understood as either viruses actively manipulating the host system to their advantage (12), or the additional draw of metabolic components for viral production simply resulting in a re-arrangement of host metabolic fluxes.

Regardless of its cause, the entanglement between host metabolism and viral production opens up the possibility to perturb the former, as a way of limiting the latter (9, 12, 13). To explore this possibility and towards understanding the potential interplay between host metabolism and the additional ‘virus demand’ on it, stoichiometric genome scale metabolic models and their optimisation through flux balance analysis (FBA) can provide ideal starting points as they are demonstrated to allow analysis of cellular physiology as an interconnected system (14, 15). Integration of virus production in a host metabolic model has already been utilised to study the infection of bacteria with phage, indicating the presence of metabolic limitations on phage replication depending on host’s metabolic environment (16). While this type of stoichiometric metabolic analysis can potentially be applied to any host-virus pair, it is particularly suited to *Alpha*- and *Flavi*-viruses. The rather simple physical and genomic structure of these viruses (17, 18) allow straightforward construction of a pseudo biochemical reaction representing their production from constituting parts. This pseudo reaction can then subsequently be incorporated into a genome-scale metabolic model of any host.

Here, we develop and apply such an FBA approach to analyse host-virus metabolic entanglement. Using a stoichiometric metabolic model of a human macrophage cell, we establish an integrated virus-macrophage metabolic model for three viruses causing current (or previously) epidemic outbreaks: Chikungunya virus (CHIKV); Dengue virus (DENV); and Zika virus (ZIKV). These are representatives of the virus genera *Alpha-* (CHIKV) and *Flavi-virus* (DENV, ZIKV), which are positive-sense single-strand RNA-viruses with rather simple physical structures (17, 18). Viruses of both families have been observed to infect monocyte derived macrophage cell lines (19-21) and are usually transmitted to humans via arthropod vectors, the most common being mosquitos of the *Aedes* genus (22, 23). By analysing the integrated metabolic model, we find that viral production results in significant alterations in host metabolic fluxes. Subsequent analysis of this integrated model through linear optimisation allows us to predict a set of host reactions, which when constrained can inhibit virus production within the macrophage metabolic system. We show that this prediction recovers most of the known targets of existing antiviral drugs, while highlighting a set of hitherto unexplored reactions that either limit virus activity broadly or for a specific virus. Thus, this computational approach allows rapid generation of experimentally testable hypotheses for novel antiviral targets within a host cell.

## RESULTS

### Host metabolism displays alternative host- and viral-optimal states

We first used the integrated virus-macrophage stoichiometric metabolic model to interrogate potential changes in host metabolism upon virus infection. To do so, we considered two idealised scenarios; one in which the metabolic system is optimised for the functional requirements of the host cell as determined by a maintenance related biomass reaction (24) (host-optimal state), and another in which the metabolic system is optimised solely for a biomass reaction that represents the production of virus particles and that is derived from viral genomes (virus-optimal state) (see *Methods* and Figure S1 for details of virus biomass calculations). These two states provide the theoretical extremes of a continuum of metabolic states that can arise during virus infection. Whilst the first scenario aims to represent the normal physiological state of macrophage cells, the second state represents a thought experiment of the host metabolic fluxes being set for maximizing virus production.

To compare the host- and virus-optimal states of the model, we analyse the metabolic fluxes directly feeding into the biomass pseudo reaction (see *Supplementary Files S1* for biomass reactions and *S2* for flux values and ranges). As expected from linear optimisation, we find that these fluxes reflect the stoichiometric differences in the amino acid and nucleotide requirements of the host cell and the virus, thus achieving perfect fulfilment of host or virus biomass requirements. We conclude that stoichiometric differences in metabolic requirements for virus production vs. host maintenance, as summarised in Figure 1, result in different metabolic flux states of the host model.

**Figure 1.**
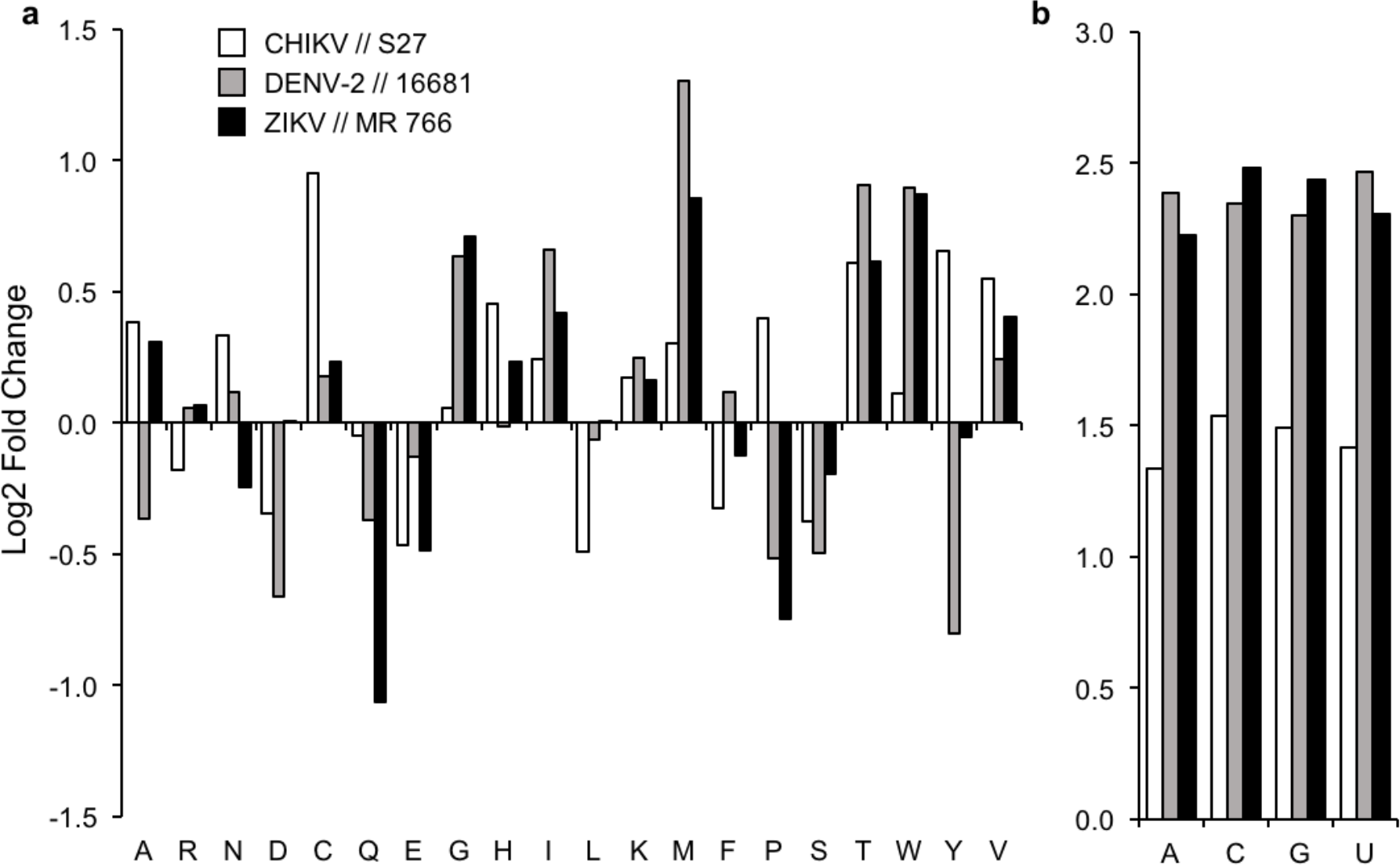
| Fold change difference in usage of amino acids and nucleotides between host and CHIKV, DENV, and ZIKV. a, b, The usage of amino acids (**a**) and nucleotides (**b**) between the host and virus biomass objective functions. The differential usage was calculated against all biomass precursors. Comparison was conducted for all 20 amino acids, and 4 RNA nucleotides (the x-axes are labelled with the common short notations for these). All calculations and biomass formulations are as described in the *Methods*, and all biomass stoichiometric values are provided as *Supplementary File S1*.

### The flux variability allowed in the system varies for viral- and host optima, indicating significant physiological changes in the host metabolism to meet viral demands

To understand how the flux changes at biomass level affect the metabolic system, we calculated the allowed flux variability for individual reactions in the model using either host- or virus-based optimisation (see *Methods*). Flux variability analysis (FVA) allows for a more robust analysis of different states of the model, compared to simply calculating optimal flux sets, which are shown to be subject to inaccuracies inherent in linear solvers used in flux balance analysis (25). We find that the median of the allowed optimal metabolic flux ranges, between host- and virus-optimal states, shows significant changes across subprocesses (Figure 2 and *Supplementary File S2*). In particular, the virus-optimal state displays significantly increased median flux for reactions associated with lipid metabolism and nucleotide biosynthesis, and significantly decreased flux for reactions associated with fatty acid biosynthesis and transport (including intracellular transport reactions). Besides these general overall trends across subprocesses, the virus-optimal state displays both increased or decreased median flux for specific reactions within each subprocess (see pie charts in Figure 2). These changes are in accordance with downstream requirements for fulfilling biomass requirements, and relate to interconnections among sub-processes. For example, the reactions from the lipid metabolism subprocess that show the most increase in their median fluxes (compared to host) involve ADP/ATP and phosphor, which are metabolites that link directly into the reactions of the nucleotide biosynthesis subprocess (and feeding into increased nucleotide requirement in the virus, see Figure 1).

**Figure 2.**
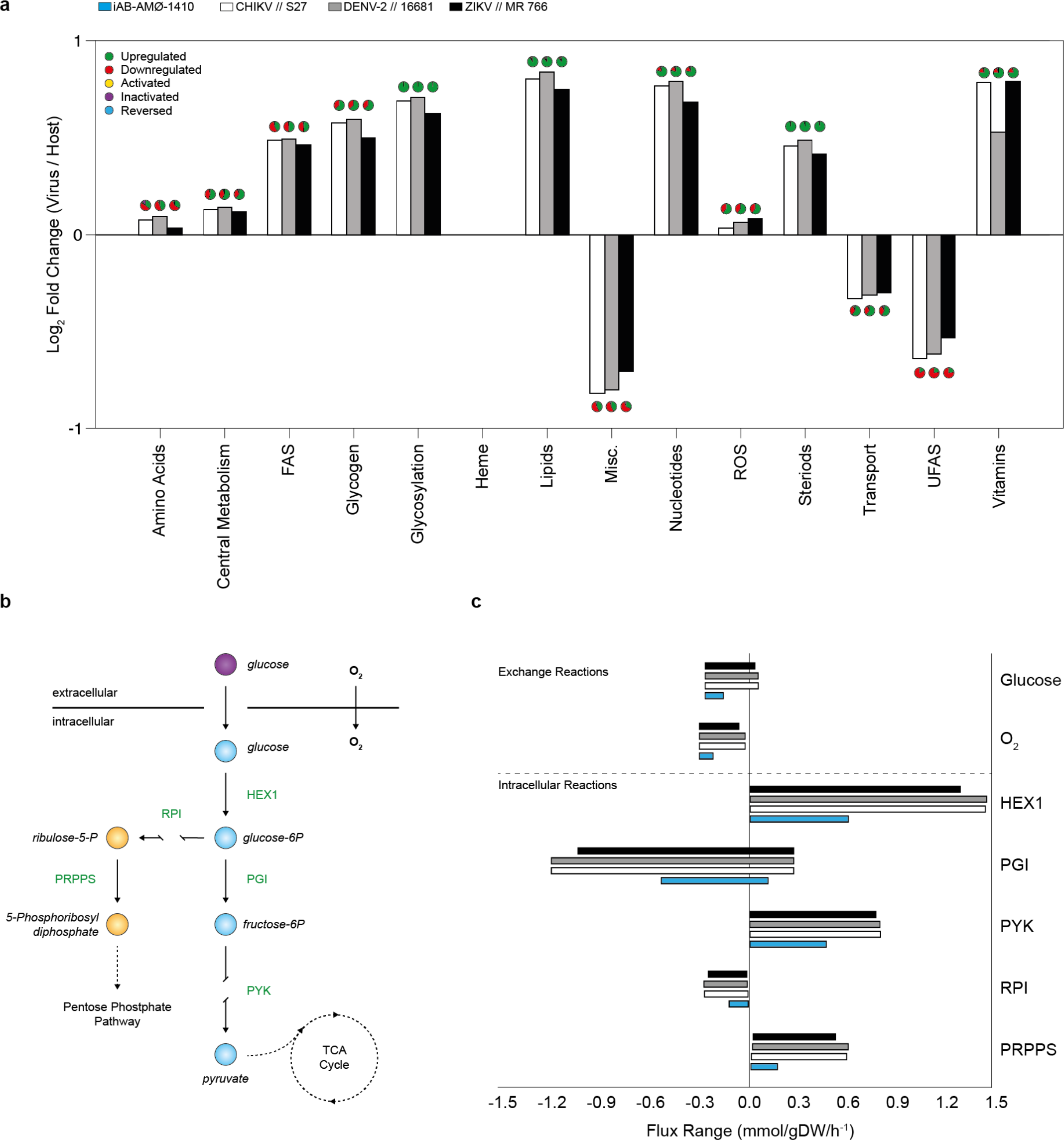
| Comparison of model fluxes between host optima and CHIKV, DENV and ZIKV optima. a, Comparisons are visualised as the sum of fluxes over aggregated subsystems using values from host- and virus-optimal states. Abbreviations used in the subsystem classification are: FAS, Fatty Acid Synthesis; ROS, Reactive Oxygen Species; UFAS, Unsaturated Fatty Acid Synthesis; Misc., Miscellaneous. The *y*-axis represents differential usage of aggregate subsystems, while the colours of the bars indicate different viruses and host (see colour coding on the panel). Positive and negative values reflect a higher or lower total flux for that subsystem in the virus-compared to host-optimal state. Pie-charts over each bar provide a summary of changes on individual reactions within a subsystem. The complete set of flux values for all reactions in the model and for all optimal states are provided as *Supplementary File S1*. **b**, Simplified schematic showing reactions involved in the glycolysis pathway. **c**, Corresponding flux ranges of individual reactions, in the glycolysis pathway, that allow attainment of host- and virus-optima. The flux ranges allowing optima for individual viruses, as well as the host are shown in differentially coloured bars with the *x*-axis showing flux values. The colour coding is as shown in panel **a**.

The specific changes in the allowed flux ranges also highlight potential physiological changes. As an illustrative example, we show the extent of changes within the glycolysis pathway, where allowed flux ranges that can sustain virus-optima are wider compared to those that can sustain host-optima (Figure 2). The allowed ranges for glucose and oxygen uptake indicate that virus-optima can be sustained even under low uptake fluxes, indicative of the potential feasibility of anaerobic metabolism still sustaining virus production (26). Taken together, this comparison of host- and virus-optimal states show that the differences within the stoichiometric requirements of the different viruses and between the host cause large-scale alterations in the host metabolic fluxes.

### Enforcing host-optimal flux ranges on individual reactions in the model predicts antiviral targets that can suppress viral production

As the host-optimal and virus-optimal flux ranges within the integrated model differ, we hypothesize that the model can be constrained in a way to limit viral production (see *Methods*). To test and utilise this hypothesis, we use the integrated stoichiometric model to identify the host reactions, which, when constrained limit virus production the most. This analysis can be implemented in different ways, for example through constraining of flux values to zero (i.e. reaction ‘knock-outs’). Applying such knock-outs, we find several reactions that limit virus-optima, but all of these also results in significant reduction in host-optima (*Supplementary File S3*). To identify if there any reactions that can perturb virus production, whilst maintaining the host viability, we constrained reaction fluxes to ranges that are derived from the FVA described above. In particular, we identified flux ranges that still allowed for the attainment of the host-optimal state, but were outside of the range allowed by the virus-optimal state (see *Methods*).

This approach highlights a set of reactions that result in different levels of reductions in the virus optima of CHIKV, DENV or ZIKV, while not affecting the host-optima (as expected from the way we set the flux constraints, see *Methods*). We identify 29 reactions that can reduce the virus optima to below 80% of the original value for at least one virus (*Supplementary File S4*). Interestingly, many of these 29 reactions are interconnected, and are involved in the *de novo* synthesis of RNA nucleotides (both purine and pyrimidine pathways) and in amino acid interconversions (Figure 3). Particular examples include reactions directly involved in the synthesis of adenosine, guanosine and uridine/cytidine nucleotides, and upstream reactions such as those involving inosine monophosphate (IMP) and orotidine monophosphate (OMP).

**Figure 3.**
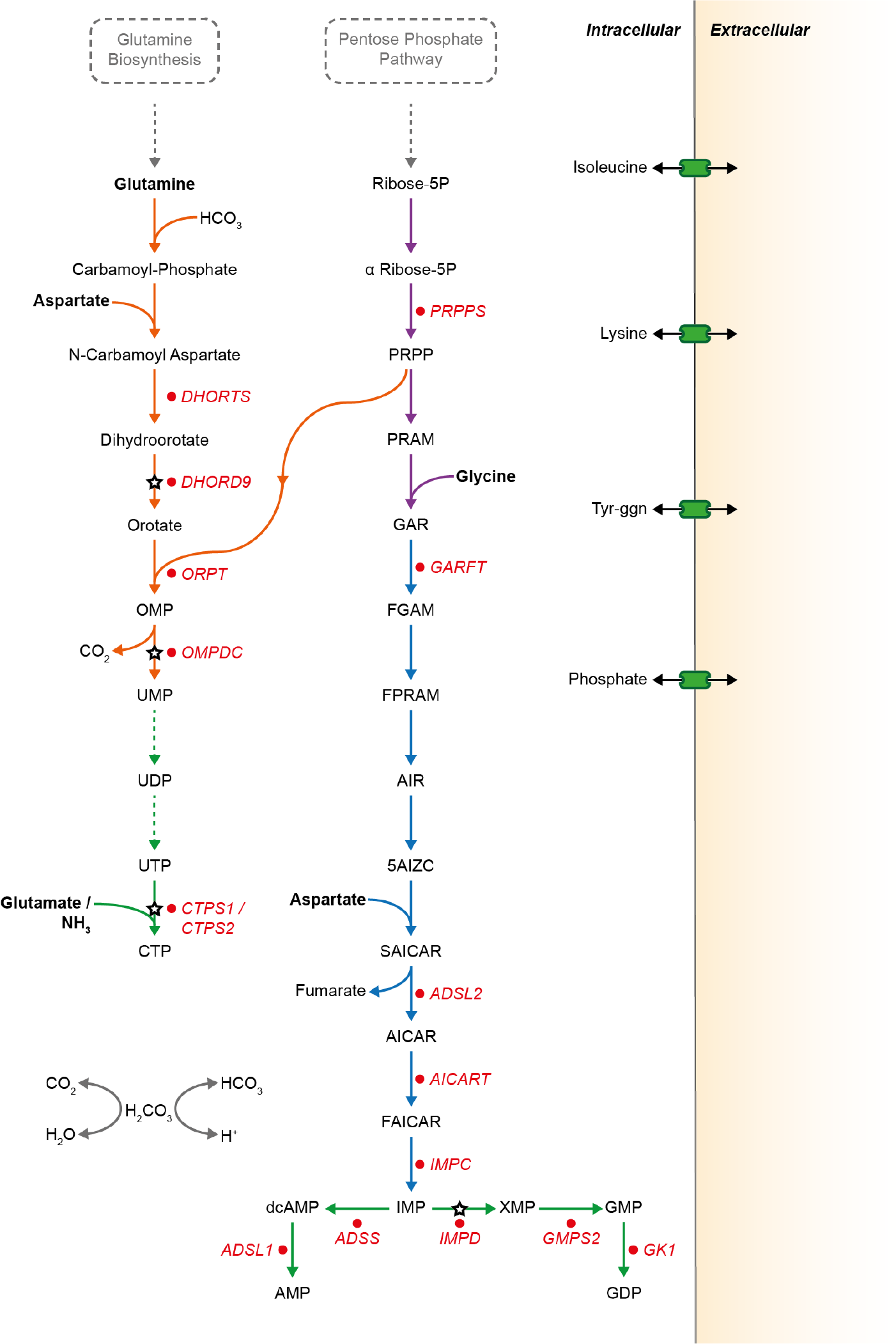
| Reaction pathway schematic showing the top 29 reactions from host-derived flux enforcement analysis and their associated antiviral drugs and inhibitors. Key reactions inhibiting virus-optima when flux ranges derived from host and virus flux variability analysis are enforced (the same reactions listed in *Supplementary File S4*). Abbreviations used for the compounds and reactions are as in *
Supplementary Table S2
* and *
S3
* respectively. Some of the identified reactions are interconnected, forming pathways.Pathways associated with subsystems are indicated by coloured reaction arrows: Orange, pyrimidine synthesis; Purple, pentose phosphate pathway; Blue, purine synthesis; Green, nucleotide biosynthesis. The starting metabolites into these pathways, Glutamine and D-Ribose 5-phosphate, are derived from glutamine biosynthesis and pentose phosphate pathways. Reactions targeted by a known antiviral or inhibitor are marked by white and red filled stars and circles, respectively. Complete list of antiviral compounds from which the matches were obtained are provided as *
Supplementary Table S1
*. Complete list of inhibitors and the associated reactions are provided as *Supplementary File S4*. A complete list of enforcement results for all reactions are provided as *Supplementary Files S2* and *S5*.

These identified reactions are potential antiviral targets, in the sense that altering their fluxes can limit virus production within the host. Thus, we explored if these reactions match with known antiviral drug targets. Performing a literature analysis, we found that there are currently 10 antivirals, specific to RNA-viruses, and these target only 5 unique metabolic enzymes (see *
Supplementary Table 1
*). Of these 5 drug targets (and the associated drugs) one has been experimentally verified to be effective against CHIKV (Inositol-5’-monophosphate dehydrogenase; *IMPD*) (27); and another against DENV (Dihydroorotate Dehydrogenase; *DHORD9*) (28). Whilst the other three targets have been verified to be effective against a number of RNA viruses (29), they are yet to be tested against CHIKV, DENV and ZIKV.

We found that out of these 5 known antiviral targets, all are implicated in our analysis. The three known antiviral target reactions involving the genes *IMPD* (27); *DHORD9* (28); and Orotidine-5’-phosphate Decarboxylase (*OMPDC*) (29) are found to perturb virus optima for all viruses (Figure 3). The antiviral target S-adenosylhomocysteine hydrolase (*AHC*) (29) is predicted to effect only CHIKV optima, and only to a level higher than the 80% cut-off we used in the above analysis (we note that setting *AHC* reaction flux to zero abolishes virus growth for all three viruses (see *Supplementary File S3*)). Finally, CTP synthase, which has been indicated to exhibit an effect on several RNA viruses (29), is included in the model as two reactions which perform the same reaction utilising either ammonia (mediated by *CTPS1*) or glutamine (mediated by *CTPS2*) as a nitrogen source (30) and therefore not highlighted in our initial flux enforcement analysis focusing on single reactions. When we constrain both reactions associated with these two reactions simultaneously at host-derived flux ranges, a reduction in all virus optima is observed.

### Identified flux enforcement effects are significant and arise from differences in host- and virus- stoichiometric requirements

The above results provide support for using integrated host-virus metabolic models for identifying host-based antiviral targets. However, any analysis based on stoichiometric metabolic flux optimisation, as done here, can be dependent on details of model implementation and assumptions (25). For example, while the host metabolic model and the biomass function that it incorporates are verified against experimental data (24), the model still assumes a specific media composition and uptake fluxes. To test if our predictions are robust against key assumptions in the model, we analysed the effects of variations in the media composition for the host model and the genomic sequence of the virus on the antiviral target prediction. In particular, we repeated the above analysis for 1000 alternative media uptake fluxes and 1000 point mutations for each of the three virus genomes, affecting the final virus biomass (see *Methods*). We found that the list of reactions with highest impact on virus biomass, while maintaining the host biomass, are qualitatively not altered with these changes in the model structure and we still recover known broad antiviral targets (*Supplementary File S5* and Figures S2-4).

To further probe the significance of these reactions as effectors of virus production, we generated 6000 randomised virus biomass compositions using the three original virus biomass functions as a starting point (see *Methods*). Repeating the enforcement analysis for this randomized set thus allowed us to generate a distribution of effects of reactions on virus-like particle production in the host system, thus acting as a null hypothesis. Results from the enforcement of the original unaltered virus sequences, as well as the point mutated virus sequences (mentioned above), were compared against the population of ‘randomised’ viruses to assess the significance of the antiviral effect (see *Methods*). This allows derivation of a significance value for the effects resulting from each reaction, when the flux enforcement analysis is applied on it. When we rank reactions according to the significance of their effects, we find that the list of reactions shown in *Supplementary File S4* are ranked among the top (*Supplementary Files S5* and *S6*), i.e. these reactions cause biomass reduction for each virus that is statistically significant when compared to their effects on randomised virus-like biomass functions. There are a couple of exceptions only in the case of ZIKV, where the reactions mediated by cystathionine g-lyase (*CYSTGL*) and cystathionine beta-synthase (*CYSTS*) did not show any significance in their effect under flux enforcement. Additional statistical analysis showed that most of the reactions listed in *Supplementary File S4* (27 out of 29) also showed significant differences in the magnitude of their effects among the three different viruses. In other words, whilst the reactions we highlight are not necessarily unique when comparing amongst CHIKV, DENV and ZIKV, their quantitative effects on virus production is significantly different for each individual species. This, combined with the fact that our randomisation process maintained the key features of stoichiometric differences among the host and virus-like biomass functions, highlight that the flux-perturbing effects of the identified reactions emerge from the core metabolic stoichiometric differences between host and the viruses. In particular, the fact that viruses use much higher levels of nucleic acids per biomass unit (Figure 1).

### Many of the predicted additional host reactions effecting virus production can be targeted by existing drugs

Considering the computationally predicted potential of the additional reactions identified as antiviral targets, we have searched for these reactions in a database of known inhibitor-like molecules (31). We found that 15 of these reactions already have known molecules, and in some cases existing drugs, targeting their catalysing enzymes (Figure 3, and full list in *Supplementary File S4*). These findings present experimentally testable predictions on host reactions, the disruption of which could limit virus production. It must be noted, however, that our computational analysis identifies flux enforcement based on differences in host- and virus-optimal states of the model, where ‘enforcement’ can mean either reduction or increase in a given flux. In contrast, most of the currently known molecules act as enzyme inhibitors (31) and would be expected to reduce metabolic fluxes.

## DISCUSSION

We present a computational approach that combines application of FBA and FVA with development of integrated host-virus metabolic models. We show that this novel approach recovers the known metabolic antiviral targets within a human macrophage cell and predicts new potential targets. These predicted reactions fall primarily onto pathways involving nucleotides and amino acids that are differentially used by the host and virus. The results of this study are in line with an integrated perspective that views the virus as an additional metabolic burden on the host cells that could be met or avoided by tinkering of host metabolic fluxes. The observed overlap between predicted reactions and known antiviral drugs gives confidence to this integrated modelling approach and highlights its potential as a rapid prediction tool to guide experimental design. This can be especially useful in the case of new and emerging viruses for which limited clinical and experimental data may be available to inform drug target identification.

The integrated stoichiometric metabolic modelling approach focuses on metabolic changes as a driver of virus production, and does not consider factors associated with virus-host cell recognition, viral entry and release (32). Furthermore, the application of the linear optimisation on stoichiometric models (i.e. FBA and FVA) strictly assumes that host metabolism is at steady state, and thus prohibits analysis of the dynamics of cellular physiology. Such dynamics could be taken into account to a certain extent by imposing different flux constraints, which could be derived from proximal experimental data (16), through development of simplified metabolic temporal models (10, 11), or by combining dynamics with linear optimisation on stoichiometric models (33, 34). Additionally, the extent of the missing information in genome-scale stoichiometric models creates limitations on how much of the metabolic processes can be covered (35).

The future efforts into model curation and standardisation (36) would open up the possibility of extensive analysis of host-virus pairings from a metabolic stance. The presented findings already suggest that targeting host metabolic processes that are linked to host-virus compositional mismatches can be used to combat virus production without altering host functions. In particular, analysis of extended flux enforcement strategies such as flux limitations on double and triple reaction combinations might identify virus specific drug combinations. Combined with the future development of additional host-virus integrated models covering many cell and virus types can thus allow a fruitful route to computational guiding of experimental antiviral drug discovery.

## METHODS

### Generation of virus biomass objective functions

To implement the FBA approach to studying virus infections from a metabolic stance, we define a pseudo reaction accounting for the production of virus particles from its constituents. We call this reaction a virus biomass objective function (VBOF). To account for metabolic fluxes associated with the virus production, the VBOF needs to capture the stoichiometry of nucleotide, amino acid and associated energy metabolites relating to virus production, similar to biomass production function used for microbial metabolic models (37). We derive the metabolic stoichiometry of virus production from the viral genome sequence, the subsequently encoded proteins, the copy number of those proteins, and knowledge of the energetic requirements for peptide bonds and phosphodiester bonds. Details of this derivation is given below, while a schematic of VBOF generation is included as Figure S1.

#### Genome and protein information for the viruses

The genome sequences used in the present study are obtained from the NCBI genomic database (38) using the following accession numbers and accessed in March 2016; Zika; NC_012532.1, Dengue; NC_001474.2, and CHIKV; NC_004162.2 (original files are provided as *Supplementary File S7*). Viruses can be classified by their replication methods, known as the Baltimore Classification System (39), and depending on this classification, a viral particle may contain more than a single copy of the genome. This is factored into the calculation of the nucleotide counts. In the presented study, all studied viruses fall into Group IV classification: they replicate their positive single-stranded RNA (+ssRNA) genome via a negative ssRNA (-ssRNA) intermediate. Therefore, the counts of the nucleotides in the negative strand is equal to the counts of the complementary nucleotide in the positive strand, i.e. count of A on (+/-) strand = count of U on (-/+) strand, and similarly for G and C counts. The count for each RNA nucleotide (Adenosine (A), Cytidine (C), Guanine (G) and Uracil (U)) can be taken directly from the virus genome sequence: RNA utilises Uracil (U) in place of Thymine (T), therefore T must be replaced with U from the genome sequence readout. In this study, all the viruses have two categories of polyproteins that compose the proteome: structural and non-structural. The amino acid sequence of these two polyproteins, and indeed any genome derived protein sequences, are obtained from gene annotations of the viral genomes as provided in the NCBI genome entries (see above for NCBI entries used). The different sub-categories of the viral proteome may be differentially incorporated into a single virus particle. For the viruses studied here, the structural and non-structural polyproteins are expressed in a ratio that is derived from the overall virus structure (i.e. proteins in the capsid, nucleocapsid, etc.)(18). The ratio is 1:240 for CHIKV(18), and 1:180 for DENV/ZIKV (17). More broadly, the ratio of different protein classes in a single virus particle can be derived from the overall virus structure or directly from literature / experimental evidence.

#### Calculating nucleotide investment per virus

The total moles of each nucleotide in a mole of virus particle (
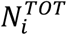
) are obtained from their count in the virus genome (
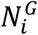
) and replication intermediates (
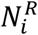
), and multiplied by the genome copy number (*C_g_
*):

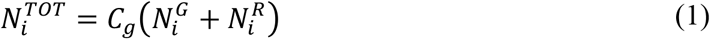

where the indexation is over nucleotides. The moles of nucleotides are then converted into grams of nucleotide per mole of virus (*g_NTPS_ mol^-1^
_virus_
*; 
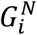
), by multiplying 
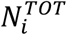
 with the respective molar mass (g mol^-1^) of the nucleotides (
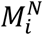
):

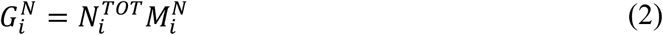

where the indexation is over nucleotides. Summing 
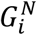
 over all nucleotides and combining this with the similar calculation for amino acids allows us to get the total molar weight of the virus in terms of nucleotides and amino acids (*M_v_
*, see Equation 15 below). Finally, the stoichiometric coefficients of each nucleotide in the VBOF are expressed as millimoles per gram of virus (*mmol_NTPS_ g^-1^
_virus_
*; 
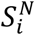
):

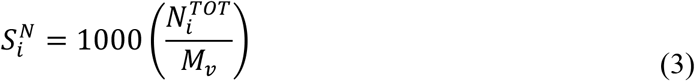

where the indexation is over nucleotides.

#### Calculating amino acid investment per virus

The total moles of each amino acid per mole of virus particle (
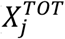
) is obtained similarly using the sequence information of structural (
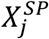
) and non-structural (
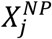
) proteins. Counts of each amino acid in these proteins is multiplied by the respective copy numbers of these proteins (*C_sp_
* and *C_np_
*):

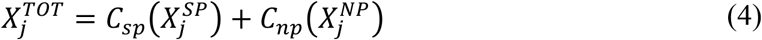

where the indexation is over amino acids. *C_np_
* is 1 for all viruses studied here, while *C_sp_
* is 240 for CHIKV (18), and 180 for DENV/ZIKV (17). The moles of amino acids per mole of virus is then converted into grams of amino acid per mole of virus (*g_AA_ mol^-1^
_virus_
*; 
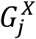
), by multiplying (
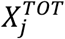
) with the respective molar mass (g mol^-1^) of each amino acid (*M^X^
*):

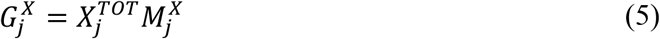

where the indexation is over amino acids. Finally, the stoichiometries of each amino acid in the VBOF is expressed as millimoles per gram of virus (*mmol_AA_ g^-1^
_virus_
*; 
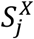
):

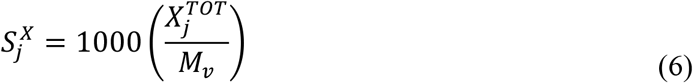

where the indexation is over amino acids.

#### Calculating ATP requirement for amino acid polymerisation (mmol g^−1^ _virus_)

The polymerisation of amino acid monomers requires approximately 4 ATP molecules per peptide bond (40), defined here as the constant *k_ATP_
* (= 4) The overall moles of ATP (*A^TOT^
*) required to form the structural (*A^SP^
*) and non-structural (*A^NP^
*) polyproteins are calculated from the respective amino acid counts:

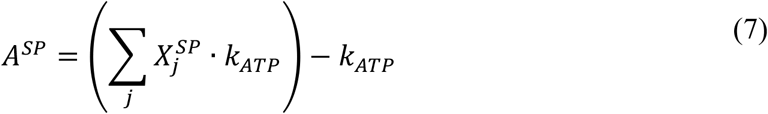

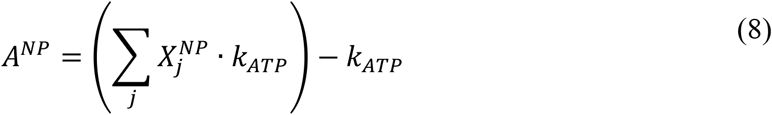

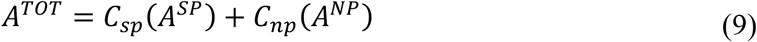

where the indexation is over amino acids. From *A^TOT^
*, we calculate the stoichiometry of ATP in the VBOF as millimoles per gram of virus (*S^ATP^
*):

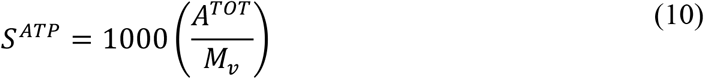

As ATP is hydrolysed in this process, the water requirement for polymerisation(
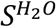
) is equal to that of ATP. The products from the hydrolysis of ATP (ADP, P_i_ and H^+^) are also accounted for in the VBOF (see Equation 16).

#### Calculating pyrophosphate (PP_i_) liberation from nucleotide polymerization(mmol g^−1^ _virus_)

The polymerisation of nucleotide monomers to form the RNA viral genome liberates a PP_i_ molecule (40), defined here as the constant *k_PPi_
* (= 1). The overall moles of PP_i_ (*P^TOT^
*) required to form the viral genome (*P^G^
*) and replication intermediates (*P^R^
*) are calculated from the respective nucleotide counts:

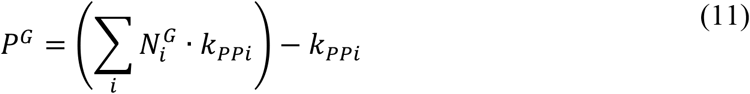

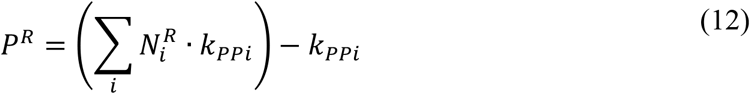

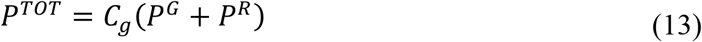

To convert this into the PP_i_ stoichiometry in the VBOF as millimoles per gram of virus (*S^PPi^
*), we again use the overall molar mass (g mol^-1^) of one mole of virus:

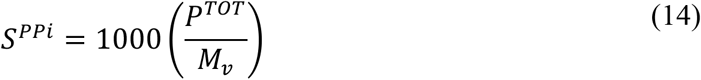

#### Calculating total Viral Molar Mass

The total molar mass of the virus *M_v_
* is calculated from the total mass of the genome and proteome components as:

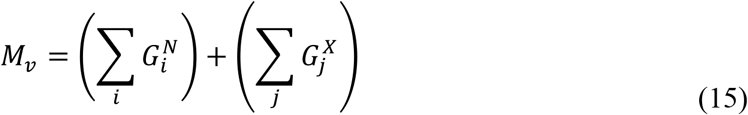

#### Final Construction of the VBOF

The left- and right-hand terms of VBOF are based on the above calculations of stoichiometric coefficients. The final stoichiometry for the VBOF (pseudo-reaction) is:

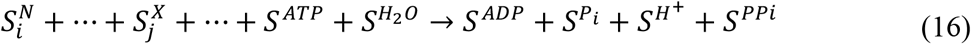

This pseudo-reaction accounts for the virus’ biomass, and the energy requirements associated with its production, and can be incorporated into stoichiometric metabolic models of the host to represent the presence of a virus in that system.

### Integration of iAB-AMØ-1410 and Chikungunya, Dengue and Zika Viruses

The VBOFs for the three viruses (CHIKV, DENV, and ZIKV) were integrated into three-separate instances of the ‘host’ macrophage model (iAB-AMØ-1410) (original files are provided as *Supplementary File S7*). In each individual case, the respective VBOF was appended into the existing macrophage model, with a lower flux bound of zero and an upper bound of infinity, reflecting no upper constraints on this flux (41). No other metabolites or reactions were added to any of the models. All of the individual flux bounds of the model reactions were used as previously set (24), but any bounds set to −1000 or 1000 are replaced with infinity, since the use of infinity, rather than arbitrarily large values, is shown to be a more robust approach to represent unbounded reactions in a linear programming model (41). We also confirmed that the use of arbitrary large bounds (such as −1000/1000) instead of infinity does not change the presented results qualitatively. A set of subprocesses, derived from known aggregate-subsystems (42), were appended as metadata to each individual host-virus model and linked with the pre-existing defined subsystems. A full description of the subsystems and mapping of reactions into these are supplied in *Supplementary File S3*. The used integrated model is provided in a computer readable (SBML) format with the publication.

### Characterising the stoichiometric differences between host and virus

For both the host (iAB-AMØ-1410) and viruses (CHIKV, DENV and ZIKV) we have pseudo-reactions that capture the metabolic requirements for the maintenance/production of their respective biomass. By comparing these pseudo-reaction stoichiometries, we can quantify the differences in amino acid and nucleotide requirements to fulfil the host or virus objectives. To do so, we calculate the fold change in nucleotide and amino acid usage by normalising their individual stoichiometric coefficients against the sum of stoichiometries of all metabolites present in the objective function (other than ATP):

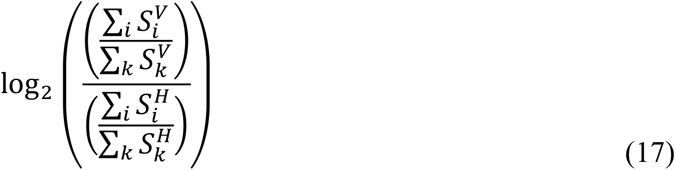

where indexation *i* is over nucleotides (or amino acids) and *k* is over all biomass precursors, and the subscript *H* and *V* indicate the use of the host and virus biomass functions respectively. A positive value indicates a higher usage for nucleotide (or amino acid) *i* by the virus than the host, whilst a negative value indicates a lower usage.

### Comparison of host- and virus-optimised states

For all analyses, the generated host-virus integrated models were optimised and reaction fluxes predicted using the linear optimisation approach known as flux balance analysis (FBA) (43). Linear optimisation is a mathematical technique that optimises a given function under a set of constraints defined by mathematical inequalities. In the context of metabolic models, the constraints correspond to limitations on reaction fluxes, while the function to be optimised can be defined as the flux in a specific reaction. While several biologically plausible objective functions can be defined (44), a common approach is to define a pseudo reaction that describes biomass production from its constituent parts, and then optimise the flux to this reaction, as we have done here. Since the set of constraints includes constraints on uptake reactions, this application of FBA results in prediction of optimal biomass production flux with respect to a specific uptake flux. In other words, FBA optimises for biomass yield from given substrates assumed to be present in the media. In this work, we apply FBA to optimise a combined host-virus metabolic system satisfy either the host or virus objective function (as described above) and study the resulting flux predictions.

To simulate a virus-optimal state, the models are optimised using the respective VBOFs of CHIKV, DENV and ZIKV viruses as the objective function, while to simulate a host-optimal state the models are optimised using the existing biomass maintenance reaction for the human macrophage as presented in (24). Besides running linear optimisation to find the optimal flux sets under each scenario, we have also performed a flux variability analysis (FVA) (45), which provides flux ranges for each reaction that still would allow attainment of a given host/virus optima. The FVA approach is shown to be more robust to instabilities associated with prediction and comparison of a single optimal flux sets (41). For each reaction in the model we compared the resulting flux ranges from FVA under host and virus optimisation, by evaluating the mean value of the allowed flux range for each individual reaction (*A_i_
*) and then collating the mean flux values for reactions associated with given subprocesses (aggregated subsystems) as a percentage of total flux through that process. More formally, we define the differential distribution of reaction flux for each subprocess (*i*) between the host and virus optimised models in terms of a fold-change:

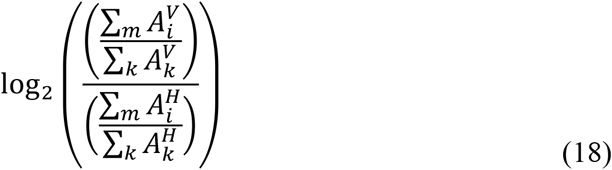

where the indexation *k* is over all reactions of the model, while the indexation *m* is over reactions that belong to subprocess *i*. The superscript indicates the use of flux values from host (*H*) and virus (*V*) optimised models respectively. A positive value indicates a higher mean flux in subprocess *i* in the virus- vs. host-optimised model, whilst a negative value indicates a lower mean flux.

### Reaction knockout and host-derived flux analyses

To find reactions which can preferentially alter virus optimised state of the model, we considered the effect of systematically constraining individual reactions. *
**Knockout analysis.**
* Knockout analysis considers the effect of systematically setting individual reaction fluxes to zero, and then attempting to maximise for VBOF. The knockout optima for the virus production reaction flux *Z_kO_
* is then compared to the original flux over this reaction; *Z_wt_
*. *
**Host-Derived Enforcement.**
* Host-derived enforcement considers the effect of maintaining a metabolic system in a host-optimised state, whilst attempting to optimise the model for VBOF. For this approach, we systematically set individual lower and upper flux bounds of individual reactions to a specific flux range. For each reaction, this range (*ε^r^
*) is derived from the corresponding minimum (*F^-^
*) and maximum (*F^+^
*) flux values for that reaction obtained from the FVA using the host (*H*) and virus (*V*) optimisation (as described above). The range (*ε^r^
*) is bounded by minimum (*ε^-^
*) and maximum (*ε^+^
*) flux values, which are given by the following conditional arguments:

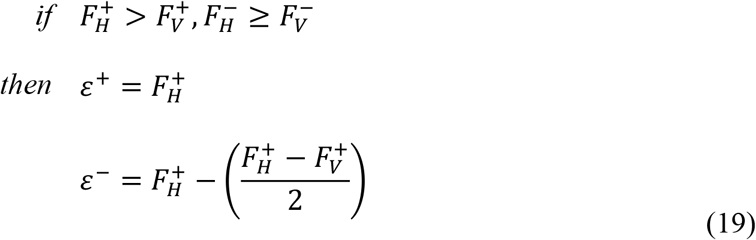

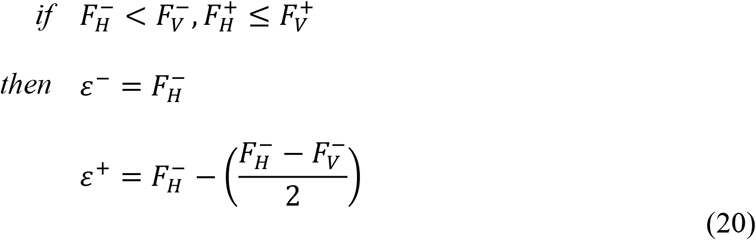

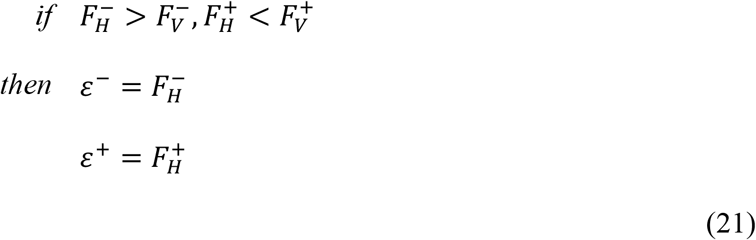

These calculated flux ranges for each individual reaction are then used to constrain the model and the model is optimised for the VBOF. The resulting optima for the virus production reaction flux; *Z*
_e_ is recorded and compared to the original optimal value; *Z_wt_
*.

### Generation of alternative virus VBOFs and statistical analyses

The presented approach evaluates the optimisation of host metabolic fluxes for virus production and how such an optimal state can be altered by specific constraints in the host model. To evaluate the impact of having different VBOFs on the outcome of such an analysis, we have generated a range of VBOFs that were either derived from the original VBOFs or were completely randomly generated. To evaluate impact of small deviations from the original VBOFs, we generated variants of the original virus genomes through nucleotide substitution; for each virus we generated 1000 genome variants, where each variant contained 1, 2, 3, 4, 5 or 10 nucleotide substitutions. For the subsequent VBOF generation from these variant genomes, the genome and protein copy numbers were kept as in the original. To evaluate more variant VBOFs, we generated another 1000 genomes for each virus that were created from the original genome with a random number (between 0 and the total length of the genome) of nucleotide substitutions, and using randomly drawn structural and non-structural polyprotein copy numbers per virus particle. Finally, and in an attempt to generate a set of VBOFs that are far removed from the original ones, both in terms of genome sequence and the structural and non-structural protein numbers, we directly generated random VBOFs. We have implemented this by drawing 1000 sets of individual stoichiometries of biomass components from a uniform distribution on [*a*, *b*], where *a* and *b* are **1)** ±99% of the original stoichiometric coefficients of a given virus, or **2)** are ±99% of the average of all original stoichiometric coefficients of a given virus. These approaches to generating variant virus genomes give us set of sequences (and associated VBOFs) that are increasingly removed from the original virus VBOFs. For each randomised VBOF created, the host-derived flux enforcement analysis was repeated (with a recalculation of the bounds used for the individual enforcements) and the reactions that perturb virus optima the most when constrained were identified. This whole analysis resulted in 8000 randomised VBOFs and FBA simulations, the results of which are summarised as percentage impact of individual host reactions on virus optima for different sets of VBOFs (see *Supplementary File S5*). To compare results of flux enforcement analysis to that obtained from using randomised biomass functions, we used a one-way ANOVA and Tukey’s honest significance tests for each individual virus against the randomised virus group, and between each individual virus species (*Supplementary File S6*).

### Measuring impact of alternative host model assumptions

As explained above, the presented analysis involves integrating a VBOF into a host model. While we used here a validated, published human macrophage metabolic model as the host model, it is desirable to evaluate the impact of specific assumptions of such a model on the results of this analysis. In particular, the macrophage model uses specific metabolite uptake fluxes, which are mostly based on experimental observations (24), but which can directly influence FBA-based results. To evaluate the potential impact of model uptake bounds, we re-analysed virus optimisation and its constraining by the host model, using alternative metabolite uptake fluxes in the host model in a systematic fashion. We first identified metabolites that are supplied (via exchange reactions) to the metabolic model with non-arbitrary lower bounds (*lb*), where *0* > *lb >-∞*. Each of the identified reactions are then systematically constrained, such that the *lb* is reduced from the original model values (24) (in steps of 10%) until the *lb* = 0, effectively knocking-out the respective exchange reaction. For each altered (additionally constrained) model, the host-derived enforcement analysis is repeated, with re-calculation of the viable host reaction bounds and optimisation of VBOF. This is done for each of the three viruses (CHIKV, DENV, ZIKV) and the results are summarised in *Supplementary File S5*.

**Supplementary Information** is linked to the online version of the paper

## Acknowledgements

This work was funded by the Defence Science and Technology Laboratory (DSTL). The funders did not have any influence in the design of experiments and analysis of the data. We acknowledge insightful discussions with Lyn O’Brien, James Findley from the DSTL and with Kalesh Shasidharan and other members of the OSS group.

## Author Contributions

OSS and SA designed and performed the research. MST involved in the original research ideas leading to this study, and both MST and AS have contributed to the interpretation of the results. All authors contributed to the writing of the manuscript.

## Author Information

The authors declare no competing financial interests. All analyses are performed using an author-developed package written in Python; the ViraNet. This analysis tool is available under an academic, non-commercial use licence on author webpages at http://osslab.lifesci.warwick.ac.uk/?pid=resources. Correspondence and requests for materials should be addressed to OSS.

## Supplementary File S1

Metabolites and their associated stoichiometric coefficients for the [host] macrophage (iAB-AMØ-1410) biomass maintenance objective function and the Chikungunya (CHIKV), Dengue (DENV) and Zika (ZIKV) virus biomass objective functions.

## Supplementary File S2

FVA results for the host-virus integrated models for CHIKV, DENV and ZIKV. Results obtained from either using the host- or virus-optimization are shown. Model reactions, model subsystems and the associated aggregated subsystems are also detailed.

## Supplementary File S3

Results for the reaction knockout and host-derived enforcement analyses. For each reaction alteration (knockout or flux enforcement), the resulting virus optima, for CHIKV, DENV and ZIKV, are shown as a percentage of the original model (without the additional constraints imposed by the analyses). The lower and upper flux bounds used for the host-derived enforcement analysis are listed for the relevant reaction and associated virus optimisation result. Model reactions and model subsystems are also listed

## Supplementary File S4

A list of model reactions identified from the host-enforcement analysis that are able to reduce the optimal flux for one of the three viruses, CHIKV, DENV and ZIKV, to below 80% of the original (obtained from the original model). Inhibitors are listed for the respective reactions, and were identified through the Brenda Enzymatic Database via the Enzyme Commission Number associated with the reaction.

## Supplementary File S5

Results of model sensitivity analysis, involving both alterations in media uptake fluxes and virus biomass functions.

## Supplementary File S6

Results of statistical analyses on the effect of host-derived enforcements across original, point-mutated, and randomly generated virus biomass functions.

## Supplementary File S7

An archived (i.e. zipped) containing: (*i*) the [host] macrophage iAB-AMØ-1410 metabolic model (‘macModel.mat’), and the (*ii*) full genbank files, obtained from NCBI, for Chikungunya (‘CHIKV.txt’), Dengue (‘DENV.txt’), and Zika (‘ZIKV.txt’) viruses. These files are used to create the virus biomass objective functions and the corresponding integrated host-virus models.

---

**Figure S1.**
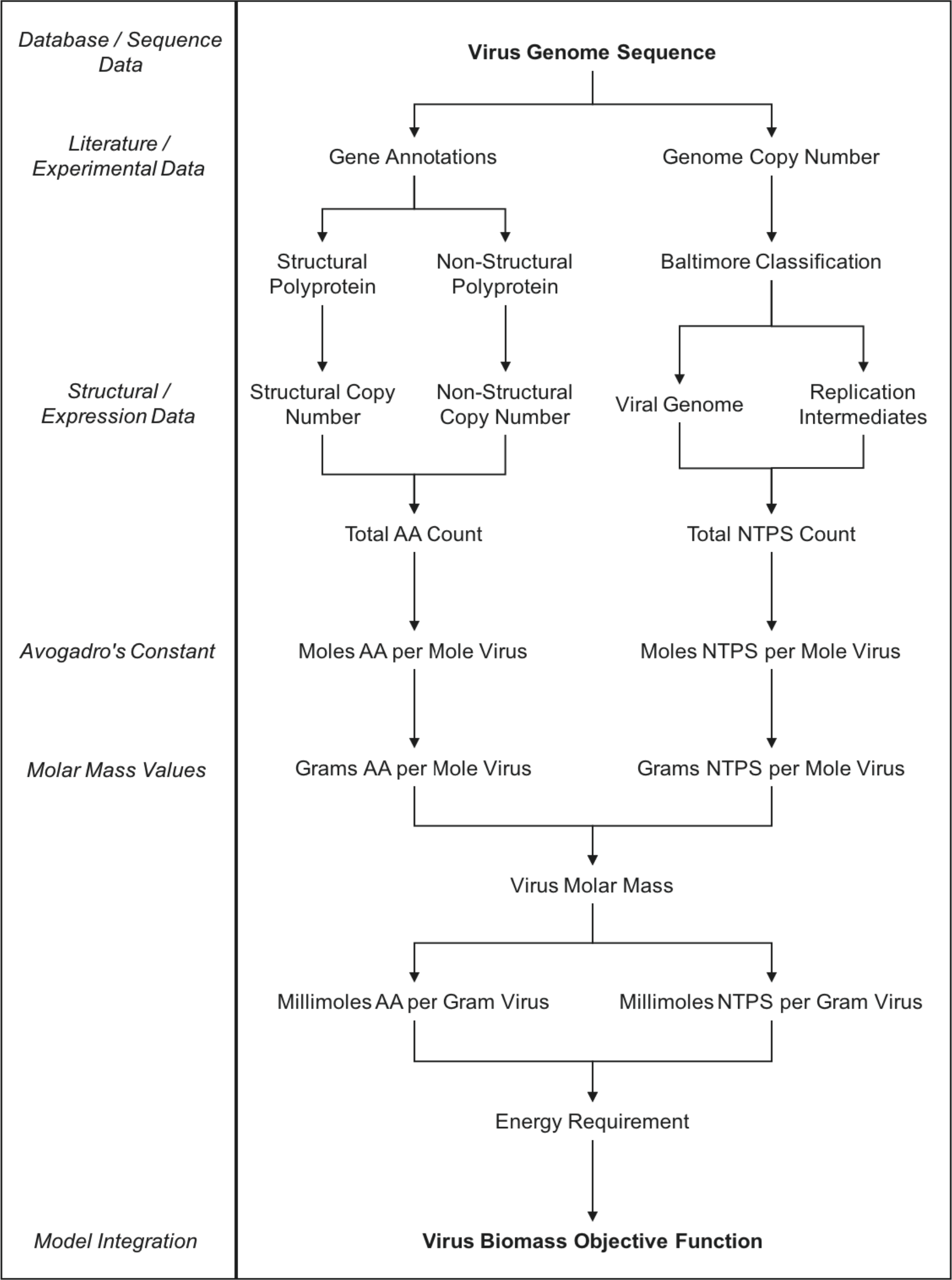
| Schematic of virus biomass objective function (VBOF) generation. Diagram outlines the process of forming the necessary components for the pseudo-reaction that represents the production of virus particles (biomass).

**Figure S2.**
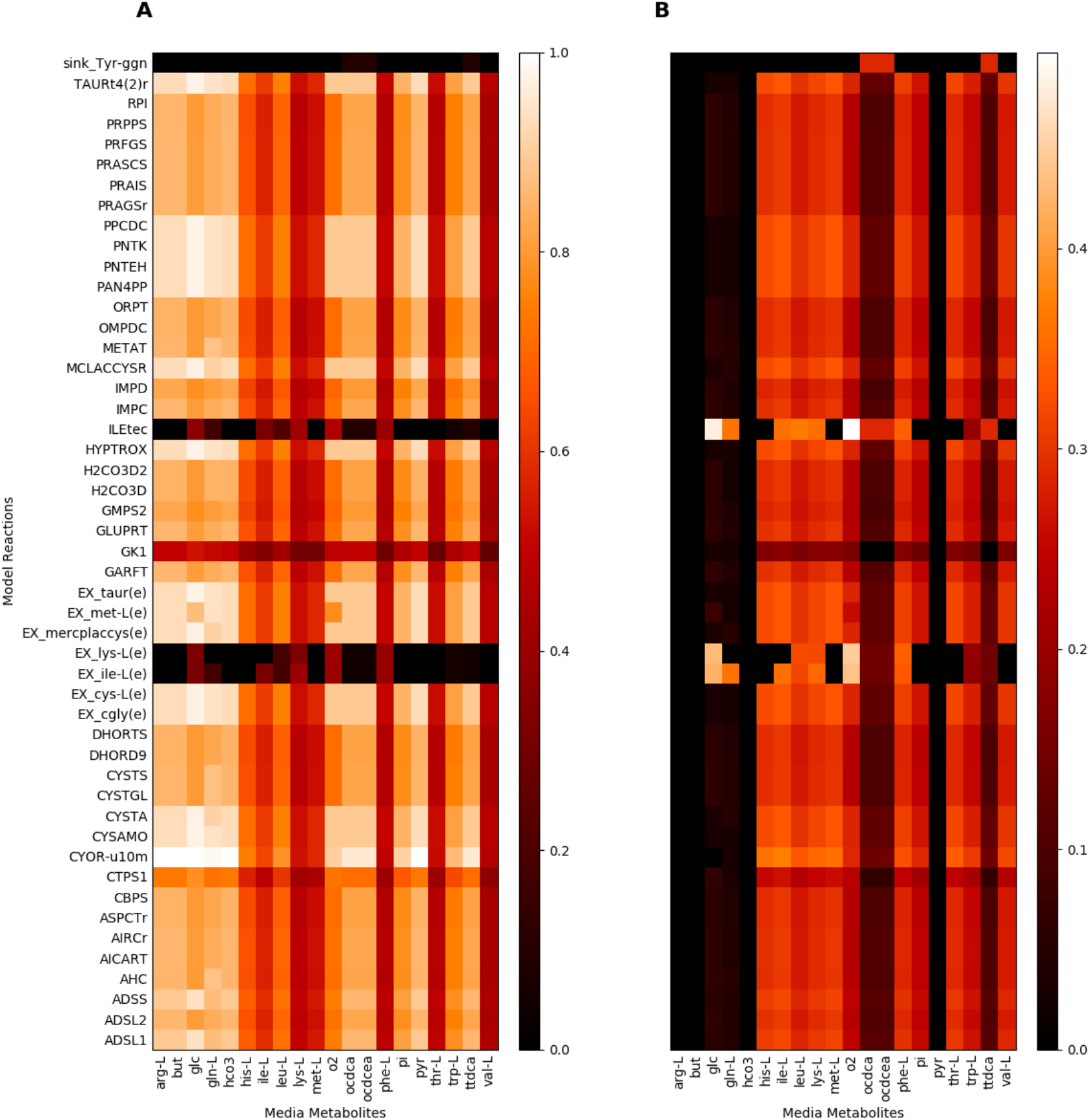
| Host-derived enforcement reactions analysed over varying model media compositions for CHIKV. A) average normalised virus optima for each media alteration condition. B) standard deviation of virus optima for each media alteration condition.

**Figure S3.**
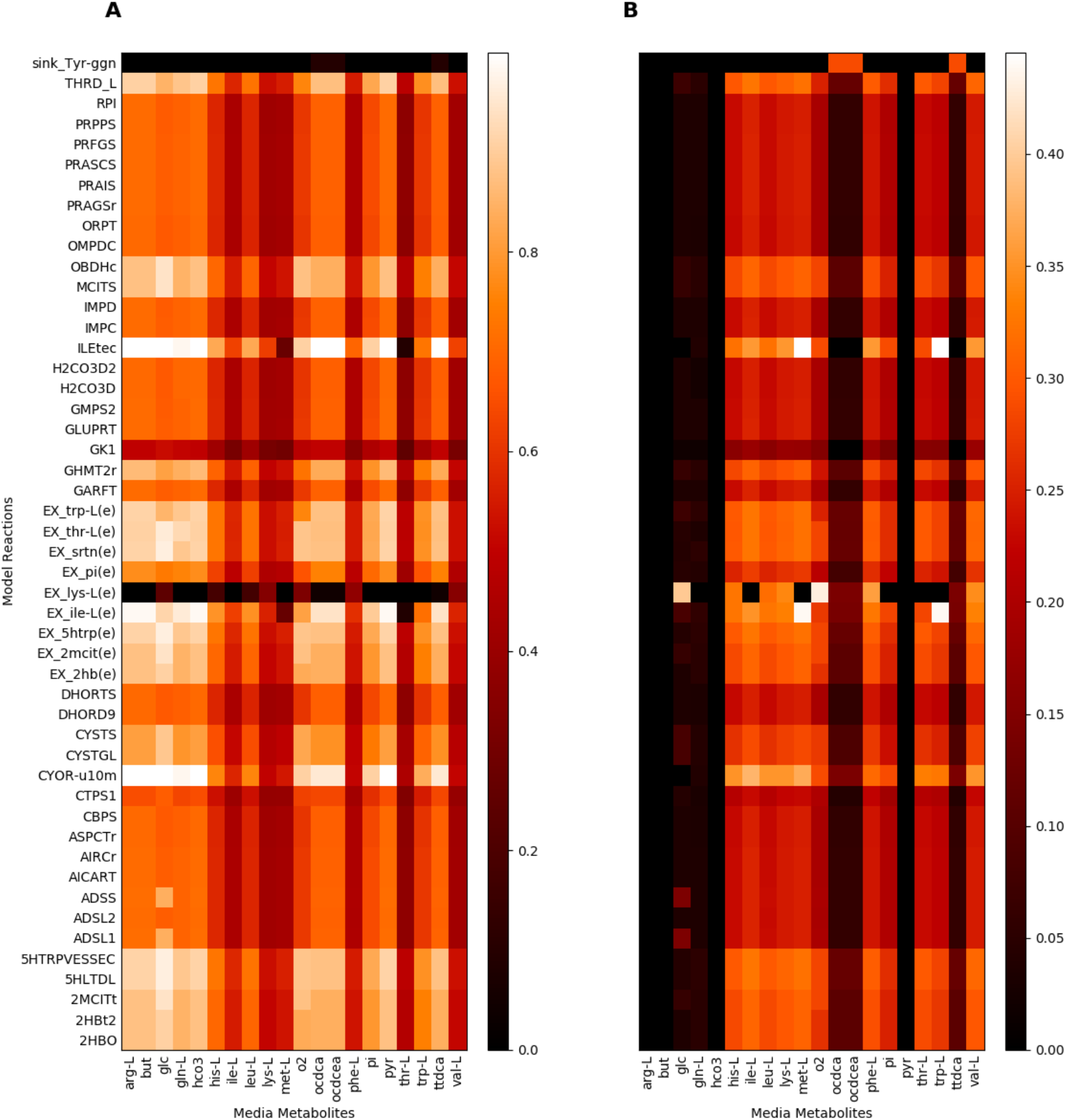
| Host-derived enforcement reactions analysed over varying model media compositions for DENV. A) average normalised virus optima for each media alteration condition. B) standard deviation of virus optima for each media alteration condition.

**Figure S4.**
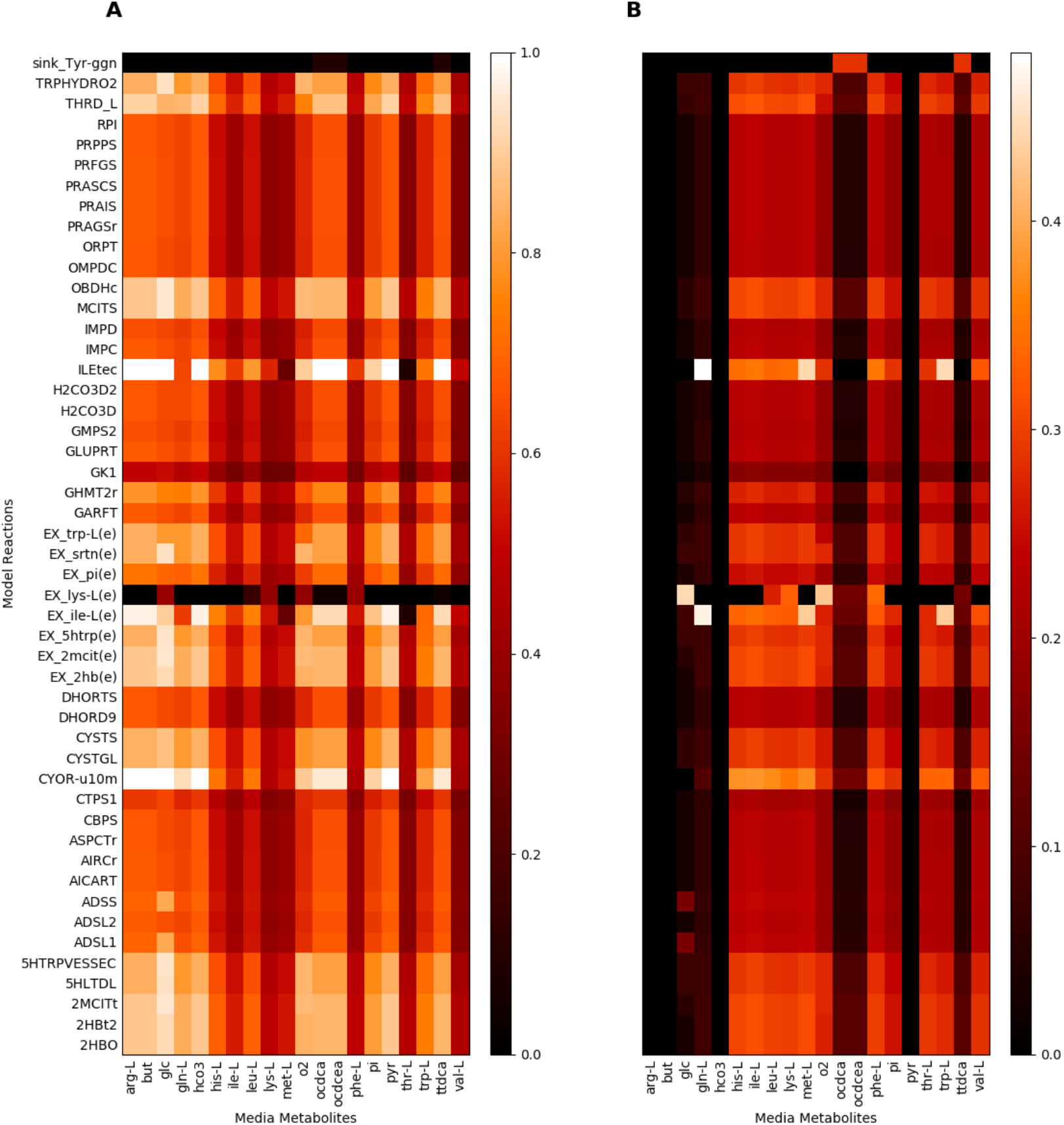
| Host-derived enforcement reactions analysed over varying model media compositions for ZIKV. A) average normalised virus optima for each media alteration condition. B) standard deviation of virus optima for each media alteration condition.

**Supplementary Table S1.**
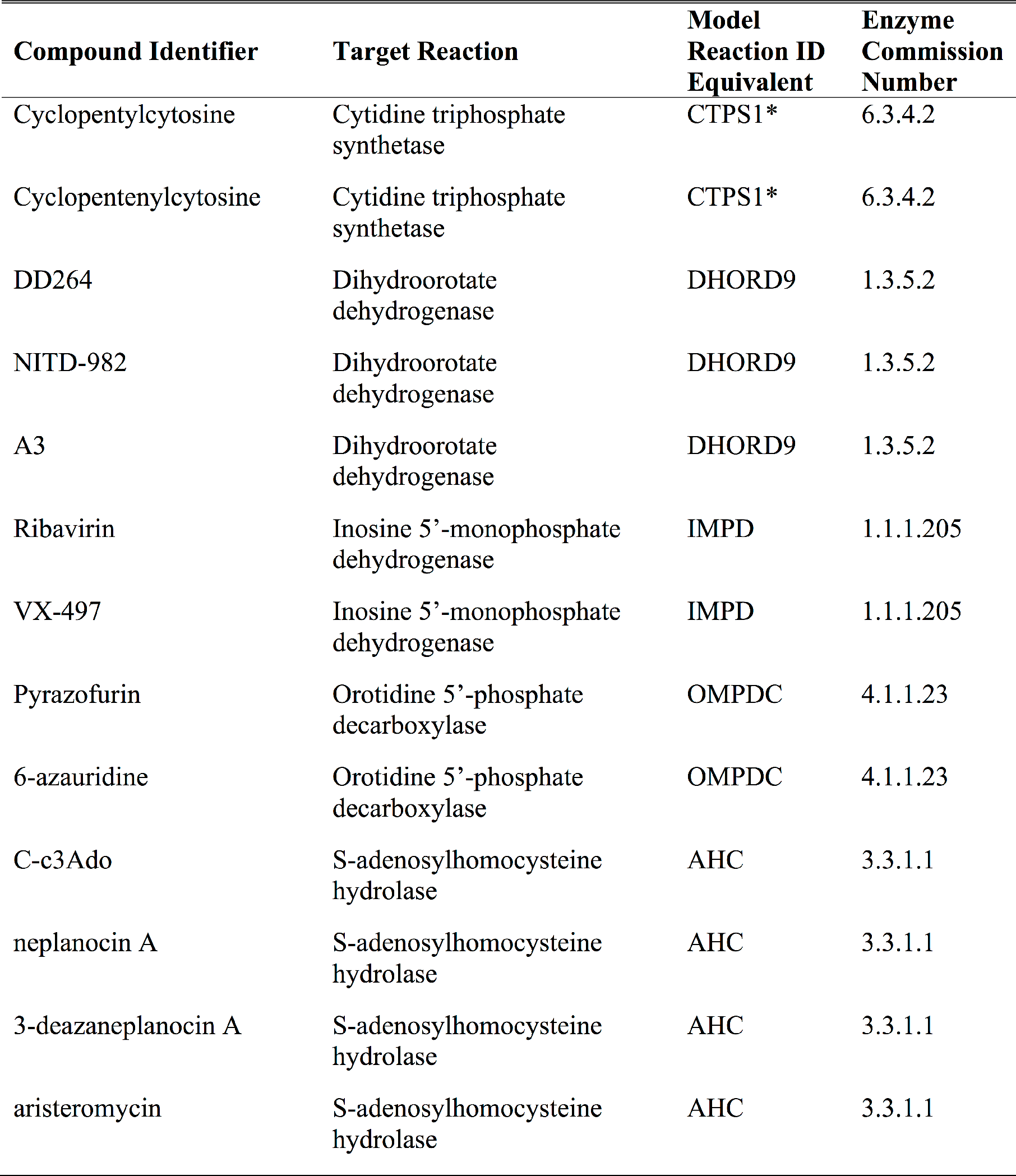
| List of antiviral compounds identified, which interact with host metabolic reactions that affect RNA virus production (29) * CTP Synthase is encoded for by two genes, CTPS1 and CTPS2, and both are affected by the associated antiviral compounds. Only one gene is shown. All antiviral compounds, and their associated target reactions, were identified from literature search (see Citations).

**Supplementary Table S2.** | Metabolite abbreviations and full names used in Figure 3, as derived from the iAB-AMØ-1410 Human Alveolar Macrophage metabolic reconstruction (24).

| Metabolite Name | Associated Abbreviation | KEGG Compound Identifier |
| --- | --- | --- |
| Orotidine 5-phosphate | OMP | C01103 |
| Uridine monophosphate | UMP | C00105 |
| Uridine diphosphate | UDP | C00015 |
| Uridine triphosphate | UTP | C00075 |
| Cytidine triphosphate | CTP | C00063 |
| 5-Phospho-alpha-D-ribose 1-diphosphate | PRPP | C00119 |
| 5-Phospho-beta-D-ribosylamine | PRAM | C03090 |
| N1-(5-Phospho-D-ribosyl)glycinamide | GAR | C03838 |
| N2-Formyl-N1-(5-phospho-D-ribosyl)glycinamide | FGAM | C04376 |
| 2-(Formamido)-N1-(5-phospho-D-ribosyl)acetamidine | FPRAM | C04640 |
| 5-amino-1-(5-phospho-D-ribosyl)imidazole | AIR | C03373 |
| 5-amino-1-(5-phospho-D-ribosyl)imidazole-4-carboxylate | 5AIZC | C04751 |
| (S)-2-[5-Amino-1-(5-phospho-D-ribosyl)imidazole-4-carboxamido]succinate | SAICAR | C04823 |
| 5-Amino-1-(5-Phospho-D-ribosyl)imidazole-4-carboxamide | AICAR | C04677 |
| 5-Formamido-1-(5-phospho-D-ribosyl)imidazole-4-carboxamide | FAICAR | C04734 |
| Inosine monophosphate | IMP | C00130 |
| Xanthosine monophosphate | XMP | C00655 |
| Guanosine monophosphate | GMP | C00144 |
| Guanosine diphosphate | GDP | C00035 |
| N6-(1,2-Dicarboxyethyl)-AMP | dcAMP | C03794 |
| Adenosine monophosphate | AMP | C00020 |
| Tyr-194 of apo-glycogenin protein (primer for glycogen synthesis) | Tyr-ggn | N/A |

**Supplementary Table S3.** | Reaction abbreviations and full names used in Figure 3, as derived from the iAB-AMØ-1410 Human Alveolar Macrophage metabolic reconstruction (24).

| Model Reaction ID | Model Reaction Name | Enzyme Commission Number |
| --- | --- | --- |
| ADSL1 | adenylosuccinate lyase (I) | 4.3.2.2 |
| ADSL2 | adenylosuccinate lyase (II) | 4.3.2.2 |
| ADSS | adenylosuccinate synthase | 6.3.4.4 |
| AICART | phosphoribosylaminoimidazolecarboxamide formyltransferase | 2.1.2.3 |
| CTPS1 | CTP synthase (NH<sub>3</sub>) | 6.3.4.2 |
| CTPS2 | CTP synthase (glutamine) | 6.3.4.2 |
| DHORD9 | dihydroorotic acid dehydrogenase (quinone10) | - |
| DHORTS | dihydroorotase | 3.5.2.3 |
| GARFT | phosphoribosylglycinamide formyltransferase | 2.1.2.2 |
| GK1 | guanylate kinase (GMP:ATP) | 2.7.4.8 |
| GMPS2 | GMP synthase | 6.3.5.2 |
| IMPC | IMP cyclohydrolase | 3.5.4.10 |
| IMPD | IMP dehydrogenase | 1.1.1.205 |
| OMPDC | orotidine-5-phosphate decarboxylase | 4.1.1.23 |
| ORPT | orotate phosphoribosyltransferase | 2.4.2.10 |
| PRPPS | phosphoribosylpyrophosphate synthetase | 2.7.6.1 |

## References

1. Littler E , Oberg B (2005) Achievements and challenges in antiviral drug discovery. Antivir Chem Chemother 16(3):155–168.

2. Mlakar J , et al. (2016) Zika Virus Associated with Microcephaly. N Engl J Med 374(10):951–958.

3. Kotzamanis K , Angulo A , Ghazal P (2015) Infection homeostasis: implications for therapeutic and immune programming of metabolism in controlling infection. Med Microbiol Immunol 204(3):395–407.

4. Maynard ND , et al. (2010) A forward-genetic screen and dynamic analysis of lambda phage host-dependencies reveals an extensive interaction network and a new anti-viral strategy. PLoS Genet 6(7):e1001017.

5. Merino-Ramos T , et al. (2015) Modification of the Host Cell Lipid Metabolism Induced by Hypolipidemic Drugs Targeting the Acetyl Coenzyme A Carboxylase Impairs West Nile Virus Replication. Antimicrob Agents Chemother 60(1):307–315.

6. Zhu Y , Yongky A , Yin J (2009) Growth of an RNA virus in single cells reveals a broad fitness distribution. Virology 385(1):39–46.

7. Yu Y , Clippinger AJ , Alwine JC (2011) Viral effects on metabolism: changes in glucose and glutamine utilization during human cytomegalovirus infection. Trends in Microbiology 19(7):360–367.

8. El-Bacha T , Menezes MMT , Azevedoe Silva MC , Sola-Penna M , Da Poian AT (2004) Mayaro virus infection alters glucose metabolism in cultured cells through activation of the enzyme 6-phosphofructo 1-kinase. Mol Cell Biochem 266(1-2):191–198.

9. Jain R , Srivastava R (2009) Metabolic investigation of host/pathogen interaction using MS2-infected Escherichia coli. BMC Syst Biol 3:121.

10. Molenaar D , van Berlo R , de Ridder D , Teusink B (2009) Shifts in growth strategies reflect tradeoffs in cellular economics. Molecular Systems Biology 5:323.

11. Weiße AY , Oyarzún DA , Danos V , Swain PS (2015) Mechanistic links between cellular trade-offs, gene expression, and growth. Proc Natl Acad Sci USA 112(9):E1038–47.

12. Maynard ND , Gutschow MV , Birch EW , Covert MW (2010) The virus as metabolic engineer. Biotechnol J 5(7):686–694.

13. Ikeda M , Kato N (2007) Modulation of host metabolism as a target of new antivirals. Adv Drug Deliv Rev 59(12):1277–1289.

14. Price ND , Papin JA , Schilling CH , Palsson BØ (2003) Genome-scale microbial in silico models: the constraints-based approach. Trends Biotechnol 21(4):163–169.

15. Terzer M , Maynard ND , Covert MW , Stelling J (2009) Genome-scale metabolic networks. Wiley Interdiscip Rev Syst Biol Med 1(3):285–297.

16. Birch EW , Ruggero NA , Covert MW (2012) Determining host metabolic limitations on viral replication via integrated modeling and experimental perturbation. PLoS Comput Biol 8(10):e1002746.

17. Mukhopadhyay S , Kuhn RJ , Rossmann MG (2005) A structural perspective of the flavivirus life cycle. Nat Rev Microbiol 3(1):13–22.

18. Strauss JH , Strauss EG (1994) The alphaviruses: gene expression, replication, and evolution. Microbiol Rev 58(4):806–562.

19. Fox JM , et al. (2015) Broadly Neutralizing Alphavirus Antibodies Bind an Epitope on E2 and Inhibit Entry and Egress. Cell 163(5):1095–1107.

20. Gollins SW , Porterfield JS (1985) Flavivirus infection enhancement in macrophages: an electron microscopic study of viral cellular entry. J Gen Virol 66 (Pt 9)(9):1969–1982.

21. Balsitis SJ , et al. (2009) Tropism of dengue virus in mice and humans defined by viral nonstructural protein 3-specific immunostaining. Am J Trop Med Hyg 80(3):416–424.

22. Garmashova N , et al. (2007) The Old World and New World alphaviruses use different virus-specific proteins for induction of transcriptional shutoff. Journal of Virology 81(5):2472–2484.

23. Lundström JO (1999) Mosquito-borne viruses in western Europe: a review. J Vector Ecol 24(1):1–39.

24. Bordbar A , Lewis NE , Schellenberger J , Palsson BØ , Jamshidi N (2010) Insight into human alveolar macrophage and M. tuberculosis interactions via metabolic reconstructions. Molecular Systems Biology 6:422.

25. Chindelevitch L , Trigg J , Regev A , Berger B (2014) An exact arithmetic toolbox for a consistent and reproducible structural analysis of metabolic network models. Nature Communications 5:4893.

26. Delgado T , et al. (2010) Induction of the Warburg effect by Kaposi’s sarcoma herpesvirus is required for the maintenance of latently infected endothelial cells. Proc Natl Acad Sci USA 107(23):10696–10701.

27. Khan M , Dhanwani R , Patro IK , Rao PVL , Parida MM (2011) Cellular IMPDH enzyme activity is a potential target for the inhibition of Chikungunya virus replication and virus induced apoptosis in cultured mammalian cells. Antiviral Research 89(1):1–8.

28. Wang Q-Y , et al. (2011) Inhibition of dengue virus through suppression of host pyrimidine biosynthesis. Journal of Virology 85(13):6548–6556.

29. Leyssen P , De Clercq E , Neyts J (2008) Molecular strategies to inhibit the replication of RNA viruses. Antiviral Research 78(1):9–25.

30. UniProt Consortium (2015) UniProt: a hub for protein information. Nucleic Acids Research 43(Database issue):D204–12.

31. Schomburg I (2004) BRENDA, the enzyme database: updates and major new developments. Nucleic Acids Research 32(90001):431D–433.

32. Timm A , Yin J (2012) Kinetics of virus production from single cells. Virology 424(1):11–17.

33. Birch EW , Udell M , Covert MW (2014) Incorporation of flexible objectives and time-linked simulation with flux balance analysis. J Theor Biol 345:12–21.

34. Mahadevan R , Edwards JS , Doyle FJ III (2002) Dynamic Flux Balance Analysis of Diauxic Growth in Escherichia coli. Biophysical Journal 83(3):1331–1340.

35. Aurich MK , Thiele I (2016) Computational Modeling of Human Metabolism and Its Application to Systems Biomedicine. Methods Mol Biol 1386(Chapter 12):253–281.

36. Zomorrodi AR , Segrè D (2016) Synthetic Ecology of Microbes: Mathematical Models and Applications. Journal of Molecular Biology 428(5):837–861.

37. Thiele I , Palsson BØ (2010) A protocol for generating a high-quality genome-scale metabolic reconstruction. Nature Protocols 5(1):93–121.

38. Geer LY , et al. (2010) The NCBI BioSystems database. Nucleic Acids Research 38(Database issue):D492–6.

39. Yu C , et al. (2013) Real time classification of viruses in 12 dimensions. PLoS ONE 8(5):e64328.

40. Haynie DT (2009) Biological Thermodynamics (Cambridge University Press, Cambridge). 2nd Ed. doi:10.1017/CBO9780511802690.

41. Kelk SM , Olivier BG , Stougie L , Bruggeman FJ (2012) Optimal flux spaces of genome-scale stoichiometric models are determined by a few subnetworks. Scientific reports. doi:10.1038/srep00580.

42. Zielinski DC , et al. (2015) Pharmacogenomic and clinical data link non-pharmacokinetic metabolic dysregulation to drug side effect pathogenesis. Nature Communications 6:7101.

43. Orth JD , Thiele I , Palsson BØ (2010) What is flux balance analysis? Nat Biotechnol 28(3):245–248.

44. Chiu H-C , Segrè D (2008) Comparative determination of biomass composition in differentially active metabolic States. Genome Inform 20:171–182.

45. Mahadevan R , Schilling CH (2003) The effects of alternate optimal solutions in constraint-based genome-scale metabolic models. Metabolic Engineering 5(4):264–276.

